# Asymmetric Binomial Statistics Explains Organelle Partitioning Variance in Cancer Cell Proliferation

**DOI:** 10.1101/2021.01.21.427596

**Authors:** Giovanna Peruzzi, Mattia Miotto, Roberta Maggio, Giancarlo Ruocco, Giorgio Gosti

**Affiliations:** Center for Life Nanoscience, Istituto Italiano di Tecnologia, Viale Regina Elena 291, 00161 Roma, Italy; Department of Physics, University of Rome ‘La Sapienza’, Piazzale Aldo Moro, 5, I00185, Rome, Italy; Department of Experimental Medicine, Sapienza University of Rome, Viale Regina Elena, 324, 00161, Rome, Italy

## Abstract

Asymmetric inheritance of organelle and cellular compounds between daughter cells impacts on the phenotypic variability and was found to be a hallmark for differentiation and rejuvenation in stem-like cells as much as a mechanism for enhancing resistance in bacteria populations. Whether the same processes take place in the context of cancer cell lines is still poorly investigated. Here, we present a method that allows the measurement of asymmetric organelle partitioning, and we use it to simultaneously measure the partitioning of three kinds of cellular elements, i.e. cytoplasm, membrane, and mitochondria in a proliferating population of human Jurkat T-cells. For this porpoise, we use multiple live cell markers which permit us both to follow the partitioning process for multiple generations and to investigate the correlations between the partitioning of different cellular constituents. Assuming a minimal model of asymmetric partitioning where cell sub-components are divided according to a biased binomial statistics, we derived exact analytical relationships for the average fluorescence intensity and its fluctuations as a function of the generation, obtaining an excellent agreement with the experimental measurements.

We found that although cell cytoplasm is divided symmetrically, mitochondria and membrane lipids are asymmetrically distributed between the two daughter cells and present a stable positive correlation with cytoplasm apportioning, which is incompatible with an independent division mechanism. Therefore, our findings show that asymmetric segregation mechanisms can also arise in cancer cell populations, and that, in this case, membrane lipids and mitochondria do not respectively segregate independently from the cytoplasm. This helps us understand the high phenotypic variability reported in these cancer cell lines. In perspective, this could be particularly relevant in the case of tumor micro-environment diversity, where comprehension of the non-genetic cell heterogeneity could pave the way to novel and more targeted therapies. Moreover, the developed experimental and theoretical apparatus can be easily generalized to different cell kinds and different cell sub-components providing a powerful tool for understanding partitioning-driven heterogeneity.

## 1 Introduction

At each division, an initial cell often referred to as the mother, firstly duplicates its genetic material and cellular component, then it halves into two daughter cells. Depending on how the internal components are partitioned, we may have a symmetric division if the daughters are identical to the mother, while in the case of asymmetric division, one or both daughter cells differentiate. In fact, while the DNA segregation process must be deterministic and exact, for other sub-components the process may be less deterministic. For this reason, it may be characterized by both a bias that determines the asymmetrical segregation towards one cell or another and noise that characterizes the segregation process stochasticity. Indeed, the molecular processes taking place inside individual cells have an inherently stochastic nature, which manifests at the level of gene expression and also in the partitioning of cellular components at the time of division^1–3^.

In certain contexts, the resulting heterogeneity may play a beneficial role in fostering differentiation and helping to promote the adaptation of cells to the environment. In populations of bacteria or cancer cells facing environmental changes, variability increases the probability that some individuals may survive the stress produced by a sudden change of the environment, e.g. the one produced by antibiotics or cancer treatments^4–9^.

Overall, while the contributions to heterogeneity coming from stochastic production/degradation processes have been widely studied^10,11^, the variability originating from partitioning is still poorly understood and often confused with the former^2,12–14^, even if, theoretical models suggest that much of the observed variability must be ascribed to noise during segregation^15^. Moreover, emerging experimental observation suggests that asymmetrical partitioning plays an important role in cell-to-cell variability, cell fate determination, cellular aging, and rejuvenation^16–18^. Whether similar mechanisms for asymmetric partitioning of cellular compounds are present also in cancer cell populations is still under scrutiny.

Semrau *et al*.^19^ and Dunn *et al*.^20^ show how pluripotent cells differentiate in progenitor and terminal cells by gradually changing the abundance of RNA and protein compounds they contain in response to the state of their gene activation network. The reason is that as several modeling approaches showed cells are complex systems in a quasi-stable state^20–24^. In this framework, cell differentiation is the effect of sufficiently large perturbations of cell compounds that bring the cell from a quasi-stable state into a new one associated with a different cell type^25^. This perspective may be considered as a mathematical complex system formulation of the Waddington epigenetic landscape conceptualization^26^.

Katajisto *et al*.^27^ have shown that mammalian epithelial stemlike immortalized cells partition mitochondria asymmetrically. Tripathi *et al*.^28^ use a computational model to show that partitioning is one of the main noise sources behind cancer cell plasticity and employs it to model epithelial-mesenchymal transition (EMT). In their model, partitioning noise increases the heterogeneity of the tumor environment and changes its ability to resist treatment^28^.

Here, we follow the proliferation of multiple populations of Jurkat cells, measuring the partitioning of different cellular compounds. To do so, we developed a method that combines a multicolor flow cytometry protocol with a computational procedure to identify cells belonging to different generations. Being a fast and non-invasive technique, flow cytometry has been successfully applied to study the growth and proliferation of very different cell types, see for instance^29,30^. Our method takes advantage of the high-throughput capability of flow cytometry to improve measurement accuracy. In particular, we analyzed the partition of cytoplasm, cellular membrane lipids, and mitochondria with the use of succinimidyl dye (CTV), lipophilic dye (PHK), and Mito-Tracker (MITO), respectively.

To explain the flow cytometry data, we use an asymmetrical binomial partitioning model, which takes into account the possibility of having biases (and thus asymmetry) in the organelle partitioning process. The asymmetrical binomial partitioning model was recently independently developed by Lijster and Åberg^31^ to model asymmetric inheritance of nanoparticles. Previously, the more restrictive symmetric binomial partition model was often adopted to model partitioning since it naturally applies binomial distributions and requires the single assumption that sub-components partition independently, and are inherited by daughter cells with equal probability^1,32–34^. This approach imposes that sub-components are segregated by a symmetrical process (symmetric segregation), and that uneven portions are the result of errors. The asymmetric partitioning model drops the equal probability assumptions and allows cellular sub-components to have a bias toward one of the daughter cells, with a single parameter *p* that measures the bias of the process and the noise. For *p* = 0.5 we have symmetric segregation and for *p* ≠ 0.5 we have asymmetric segregation. A recent work of Lijster and Åberg^31^ discusses the fact that to estimate the asymmetry from the fluorescence or particle count distribution we can not use the mean but propose to use the coefficient of variance of the full population. Unfortunately, the coefficient of variation is dependent on the relative frequencies of the generations of the cells in the population and for this reason it changes in time dynamically. With our analytical discussion, we show that under the asymmetric partitioning model assumptions, the variance of any single generation is constant. Furthermore, if we compute the variance of the cells in each generation, we can measure the asymmetry without taking into account the time of the measurements.

Moreover, for the first time to our knowledge, our multi-label method also enables us to investigate the correlation between the organelle partitioning. In fact, cellular partitioning involves several molecular compounds with different sizes and spatial distribution. Correlations at the level of the partitioning may then promote the experimentally observed nontrivial correlations at the phenotypic level^13,35–37^.

## 2 Measuring the organelle partitioning

To quantitatively measure the segregation of the cellular sub-components, we devised a multicolor flow cytometry experiment. Flow cytometry is a non-disruptive technique that enables the simultaneous acquisition of thousands of cells per second. The main idea of a multicolor flow cytometry approach is to have an initial population with two or more cell compartments stained with a live cell marker that does not affect the cell cycle and the partitioning process. The opportunity to stain more than one compartment in live cells allows us to follow how these compartments are partitioned in time and to study the guiding dynamical processes.

In our experimental setup, we used three fluorescent dyes, CTV, PKH, and Mito-tracker, which respectively label the cytoplasm, the cell membrane, and the mitochondria (see Materials and Methods for details). In particular, Métivier *et al*.^38^ used MitoTracker Green FM to assess mitochondria content in cells. This dye was described in flow cytometry protocols to establish mitochondria amount in human lymphoblastic cell lines^39^. It is of relevance that to our knowledge it is the first time that MitoTracker is used in sorting experiments to follow mitochondria partitioning in mammalian cell progeny.

After staining the population with the fluorescence dyes, we selected an initial population with a narrowly peaked fluorescence distribution. In fact, as all the cells of each generation divide into two daughter cells, the fluorescence distribution decreases its mean of one-half, and changes its variance as a consequence of the partitioning events; so the narrower the initial population the easier it is to discriminate the subsequent generations. To generate a narrowly peaked fluorescence distribution, we used fluorescence-activated cell sorting (FACS) that allows isolation of a given cell population based on the fluorescence signal of a specific color, channel, subtracted by the background signal. Once established a gate of interest, based on each signal fluorescence intensity upon acquisition, all the cells enclosed in that gate are collected in vital and functional conditions. Isolating the cells with the sorter ensures highly purified and sterile cells that can still be grown in culture for up to three weeks.

The human cancer cell line we used, Jurkat T-cells, is commonly used for proliferation studies, and it has been verified especially for multiple dye staining and labeling efficiency, showing low toxicity^40^. Indeed, in each sorting experiments we performed, cells stained for live-dead compounds were highly viable (see Figure S1 in the Supporting Information about the sorting gate strategy of the dead-exclusion diagram). Importantly, cells were passaged only twice upon defrosting to avoid excessive exposure to culture conditions prior to the experiment.

In Figure 1a, we show a schematic representation of the experimental setup. Figure 1b displays how cells are collected by sorting the cells with a high intensity signal of PKH and CTV as an example and then put in culture (Figure 1c). In many proliferation analyses, there is a tendency to synchronize cells using serum starvation to avoid that cells are in different stages of the cell cycle or temporally separated by a minimum of one division round. On the other hand, it has been found that Jurkat cells can be resistant to serum deprivation, and consequently that synchronization could be a source of error^41,42^. For this reason, we decided to not use whole-culture synchronized cells since we wanted to analyze cells in their proliferation state without perturbing the partitioning process from the outside experimentally. Furthermore, a recent work by Cooper^43^ discusses the issue of cell population synchronization, arguing that whole-culture synchronization methods are unreliable because in most cases they relay on stopping the cell cycle at a certain point, and inhibiting cell division. This procedure would lead to biases in cell phenotype, for example whole-culture synchronized cells exhibit broader cell size distribution in comparison to the unperturbed cell at the same cell cycle point. Unexpectedly, narrow sorting of stained cells may instead be effectively used to obtain unbiased cell synchronization. The fact that narrow fluorescent sorting indirectly produces cell synchronization is confirmed by two fundamental observable characteristics of a unbiased synchronization method which are cells with a narrow-size distribution, and clear picks corresponding to successive generations emerging an disappearing in correspondence of the inter-division time^43^. In our experiment, from the forward and side scattering data we observe a narrow-size distribution compared to the unsorted cells size distribution (see SI). Furthermore, even if we do not sample several time points we find that after each successive inter-division time period, a new peak (associated to a new generation) emerges and contains the near total majority of the cells of that generation. These observations confirm us that the populations were indirectly synchronized by the narrow gate on the florescence intensity. However, it is important to point out that the synchronization is a result of the sorting step, but in general our method does not require synchronization to measure asymmetry since can track generations via the fluorescence intensity.

**Figure 1.**
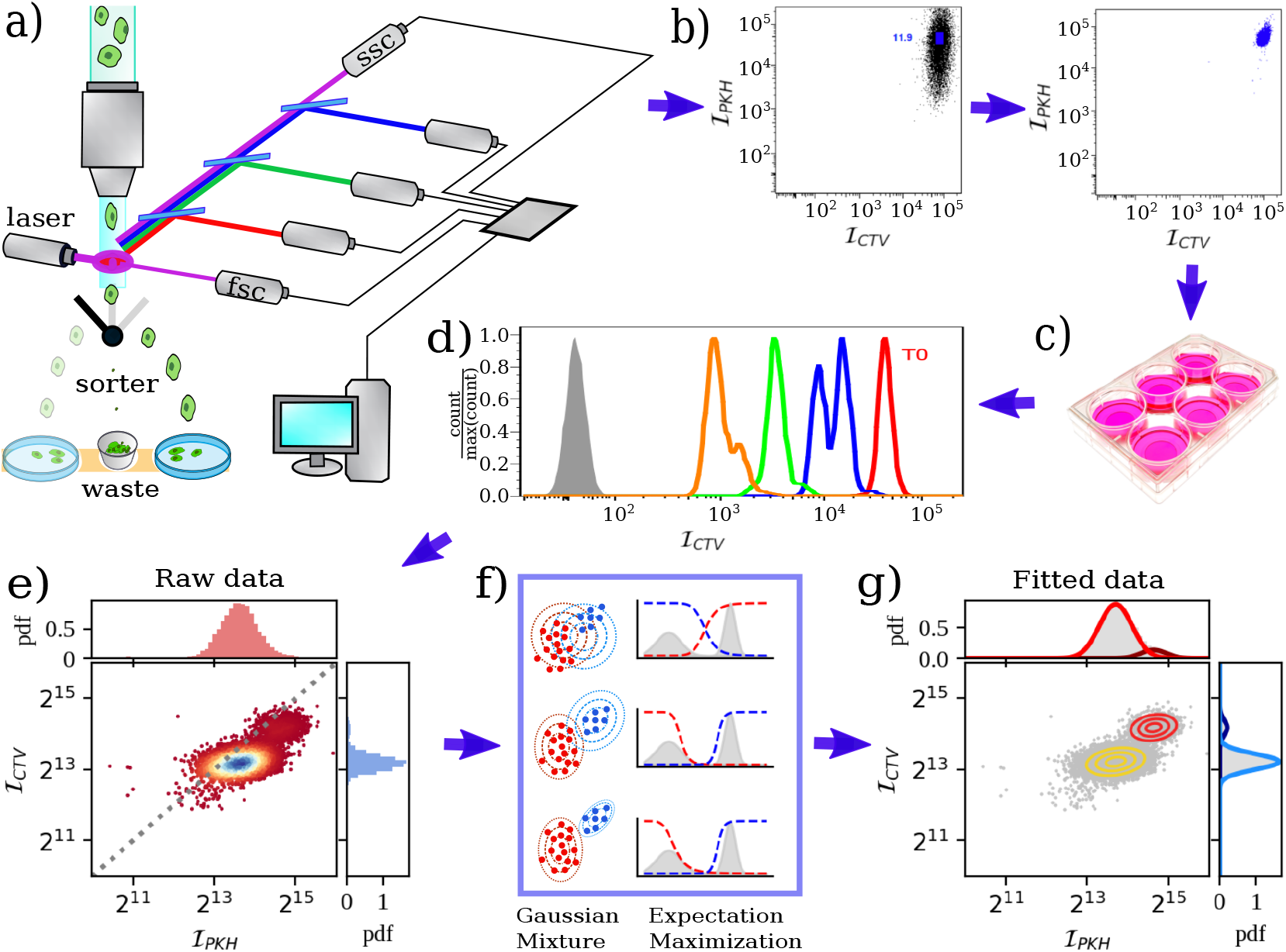
Experimental protocol. **a)** Schematic representation of a fluorescence-activated cell sorter (FACS) working principle. Stained cells pass one at a time in front of a laser source that interacts with the fluorophores of the markers. The emitted light is collected and analyzed. Eventually, a sorter divides cells according to certain thresholds on the measured intensities. **b)** The dot plot shows how cells can be collected on the basis of the highest expression level of PKH and CTV. The left panel presents a pre-sorting sample with the established collection gate and the right panel shows a post-sorted sample. **c)** Sorted cells are then put in culture for subsequent acquisition of aliquots at the flow cytometer at different time points. **d)** The histogram shows the CTV fluorescence intensity distribution that starting from the highest level at time zero (*T*_0_, red) decreases in time as each division occurs (blue, green, and orange). The grey filled distribution represents the background intensity. **e)** Two-marker dot plot. Each dot represents the measured intensity of PKH and CTV marker in a single cell. **f)** Analyzing the data with the EM mixture of Gaussians algorithm enables us to identify different populations/generations.**g)** Same as in e) but with cells clustered by generations according to the Gaussian mixture algorithm.

In practice, once the initial population is characterized, we start the measurement of the population dynamics by sampling a small portion of the growing population at an established kinetic. Then we estimate the fluorescence distribution of the population from the sampled distribution (see Methods). This enables us to follow how the fluorescence decreases at each analyzed time point (Figure 1d). The simultaneous measurement of the fluorescence intensity for all the dyes enables us to associate a 2 or 3 dimensional point for each sampled cell, **f** = (*f, g, h*), where each dimension is the intensity of one of the fluorescent markers used. The fluorescent sub-components are divided into the new daughter cells within each new generation of cells; thus, at each division the average fluorescence of the new population will be half of the previous. Consequently, if we plot the base-two logarithm of **f** on a scatter plot, and if the initial variance of the fluorescence intensity of the cells at the first generation **f_g=1_** is made sufficiently narrow (see Fig. 1b), we find that the different generations of cells form clusters on the scatter plot, and peaks on the histograms of the fluorescence intensities. Moreover, if we plot the histogram of the base-two logarithm of either *f, g* or *h*, we see that each peak is at a unit distance from the next. In conclusion, if the initial variance of **f_g=0_** is sufficiently small, Var[**f_1_**] << E[**f_1_**], we get a sequence of peaks one for each generation *g* as shown in Fig. 1d.

Each snapshot can be described as a mixture distribution,

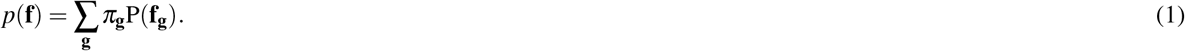

Each peak would have an expected value such log_2_(E[**f_g+1_**]) = log_**2**_(E[**f_g_**]) – 1. If we assume that P(**f_g_**) is approximately Gaussian we can model the data as a mixture of Gaussians

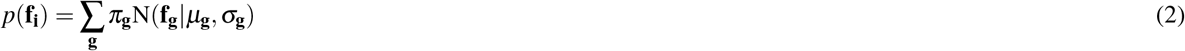

where N(**f_g_**|*μ*_g_, *σ*_g_) is a Gaussian distribution with mean *μ_g_* and variance *σ_g_* (see Materials and Methods)

Figure 1e-g shows the mixture of Gaussians protocol that assigns to each point in the clouds to the corresponding generation. We get sequences of samples that produce a snapshot of the population distribution at different times, by performing measurements at regular intervals during the proliferation. Figure 2a) shows an example of the population proliferation, followed through the intensities of the CTV and PKH markers. In this way, the sampled cells produce a series of clouds composed of data points, each of which represents a different generation. Each dot of these clouds corresponds to a cell that was part of the sample. One can spot the succession of peaks associated with the different generations that succeed one after the other in the population that grows and proliferates over time.

**Figure 2.**
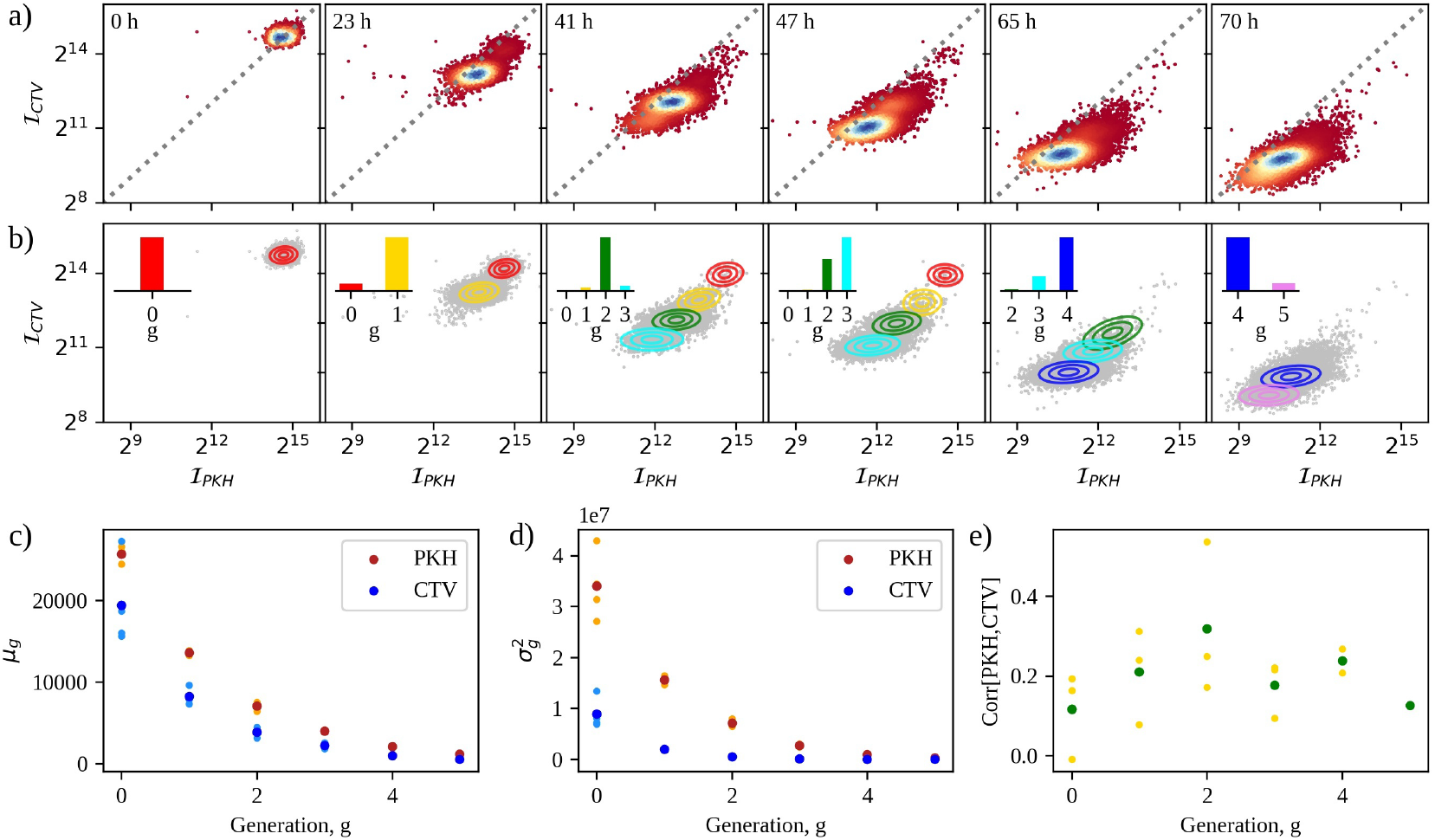
Data analysis. **a)** Dot plot of the intensity of PKH vs CTV, measured at different times (hours) in one of the three experiments with simultaneous PKH and CTV markers. **b)** Gaussian mixture fitt from the estimate model parameters obtained from a). Different colors correspond to different generations. The percentages of cells in each generation are reported in the insert as bar plot. **c)** Mean value of the intensity of the fluorescent markers as a function of the generations, obtained through the Gaussian Mixture fitting. Dark blue and red dots correspond to the mean of the intensity for each generation. **d)** Same as in c) but with the square root of the intensity variance. **e)** Pearson correlations between the PKH and CTV intensities as a function of the generations. Green dots represent the mean correlation for each generation.

Applying the Gaussian Mixture fitting procedure to each snapshot (see Figure 2b), we can measure the mean (*μ*_g_) and the variance (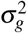) of the peaks corresponding to different generations. As we can see from Figure 2c-d both the mean intensity and its fluctuations decrease as a function of the generations for the two displayed markers. The same behavior is also present when considering the mitochondria marker (not shown in the figure). On the other hand, the Pearson correlation between the two intensities appears to remain constant during the proliferation, as shown in Figure 2e.

## 3 Model

### 3.1 Binomial partitioning

To acquire quantitative insights on the partitioning mechanisms responsible for the observed dispersion of fluorescence intensities, we defined an asymmetric binomial partitioning model. This model is a stochastic model that enables us to derive the parameters of the partitioning processes from experimentally measured means and variances of the intensity peaks. In the asymmetric binomial partitioning model, we assume that cell sub-components segregate randomly according to a Bernoulli process. To uniquely label all the cells in a progeny derived from a single progenitor cell with label 1, we consider that each cell *i* divides into two daughter cells that we can label unambiguously as 2*i* and 2*i* + 1. After *g* generations, the cell lineage generated from the cell 1 forms a lineage tree. If we start with one cell, at generation *g* = 0, each generation is composed of *n_g_* = 2*^g^* cells (Fig. 3a).

**Figure 3.**
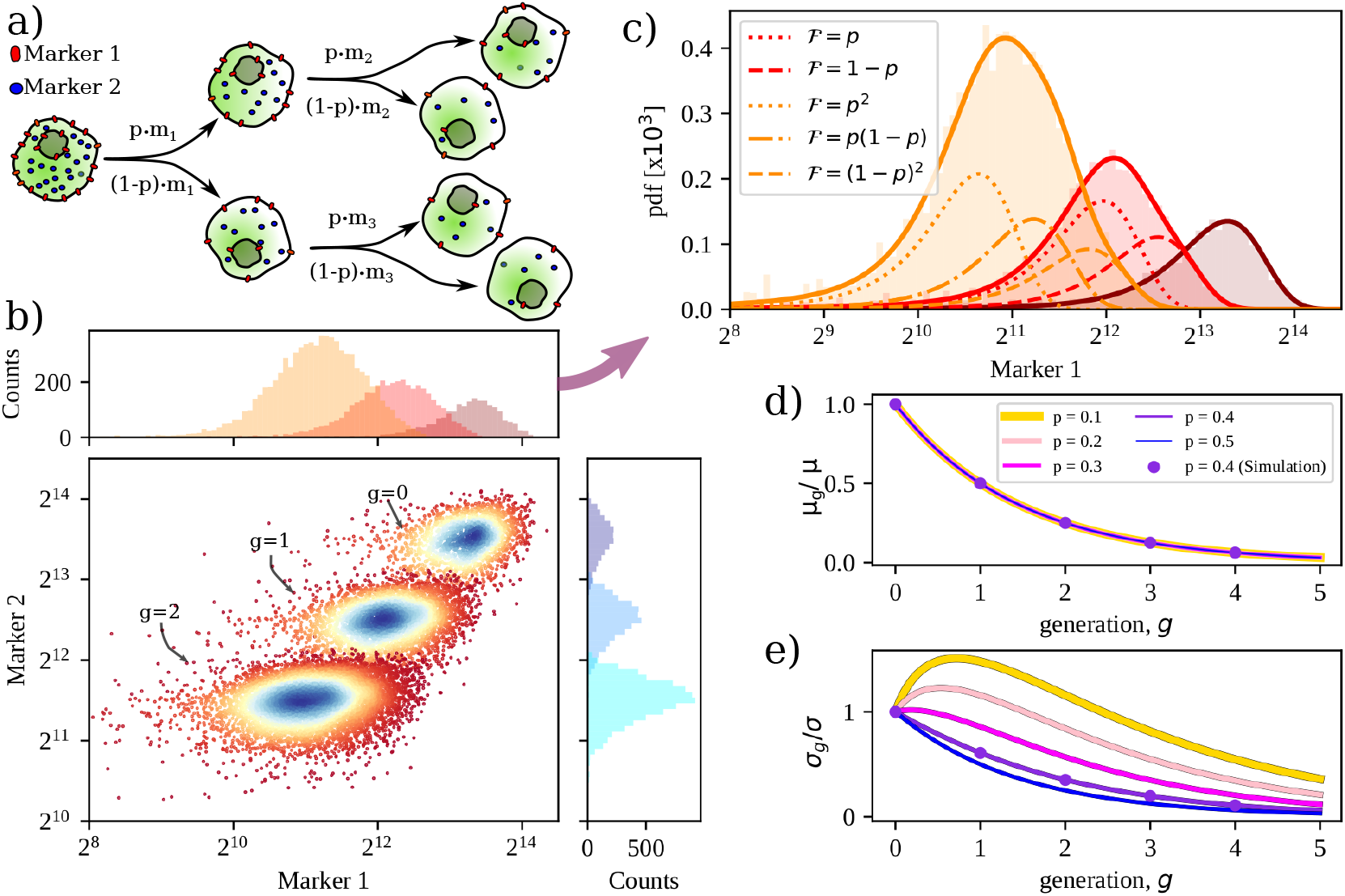
Theoretical predictions for the Bernoulli partition process. **a)** Partitioning noise and asymmetry during proliferation. When a cell *i* with *m_i_* sub-components divides, *m*_2*i*_ sub-components go to one daughter cell and *m*_2*i*+1_ = *m_i_* – *m*_2*i*_ sub-components go to the other daughter cell. **b)** Number of marker 1 and 2 per cells. Each cloud of points corresponds to a different generation. Histograms represent the marginal distributions for the two markers. **c)** Marginal distribution of marker 1 copy-numbers. For each generation, the dotted curves are associated to the different daughter cells sub-populations that originated from the previous generation. **d)** Mean number of marked compound per cell, *m_g_* as a function of the generation, g for different possible values of the partition fraction, *p*. Solutions of Eq. 17 are represented with lines while dots stand for simulated values. **e)** Square root variance of the number of marked compounds per cell, *σ_g_*, as a function of the generation, *g* for different possible values of the partition fraction, *p*.

Given that the mother cell, *i* has *m_i_* sub-components that are stained with a fluorescent dye, the daughter cells 2*i* and 2*i* + 1 will have respectively *m*_2*i*_ and *m*_2*i*+1_ (with *m*_1_ = *m*_2*i*_ – *m*_2*i*+1_) fluorescent sub-components. If we assume that the segregation is symmetric and noiseless, the cell sub-components would always be divided exactly in half, i.e. *m*_2*i*_ = *m*_2*i*+1_ = *m_i_*/2. Otherwise, if we assume that each component selects cell 2*i* or 2*i* + 1 with probability *p*, we find that *m*_2*i*_ and *m*_2*i*+1_ are random variables and that, given *m_i_*, the distribution of *m*_2*i*_ is binomial

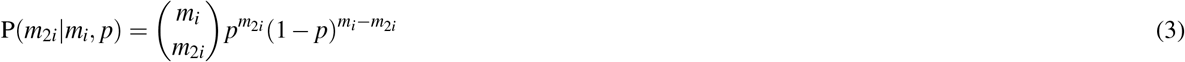

with mean *p* · *m_i_* and variance *p*(1 – *p*)*m_i_* as depicted in Figure 3a. In our flow cytometry experiment, at generation zero, we do not have a single mother cell but a population of cells characterized by a distribution over possible values of *m*_1_, P(*m*_1_), which has a specific average *μ* and variance *σ*^2^. The probability distributions of the inherited fluorescent sub-components in the first generation of daughter cells is given by the superposition of two sub-populations formed by the daughter cells 2 and 3 with respectively P(*m*_2_) and P(*m*_3_),

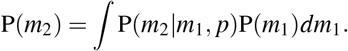

and

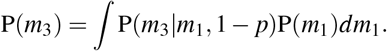

In general, given a cell *i*, and given the number of fluorescent sub-components P(*m_i_*), the distributions of the daughter cells 2*i* and 2*i* + 1 are

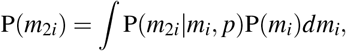

and

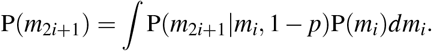

#### 3.1.1 Symmetric case

To derive a closed-form equation for the expectation and the variance of P(*m_i_*), we start with the simpler symmetric partition case (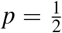), where P(*m*_2*i*_|*m_i_*) and P(*m*_2*i*+1_|*m_i_*) have average 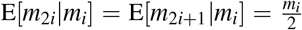 and variance 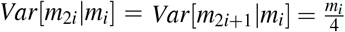.

We begin with the expression of the expectation using the total expectation law, which is just a consequence of the double-integrals computed on the expectation value. Thus, setting *μ*_1_ = *μ*,

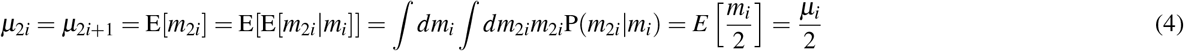

For the variance, the computation is a little more complex, but analogously, from the law of total variance, which is itself a consequence of total expectation law, and given *σ*_1_ = *σ*,

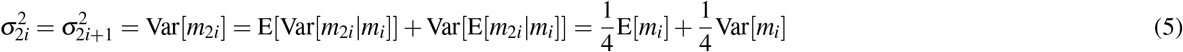

For *i* = 1,

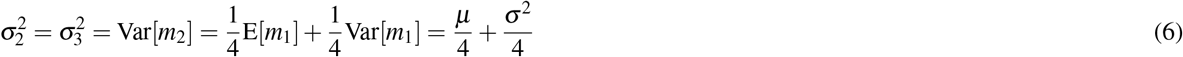

and for *i* = 1,2

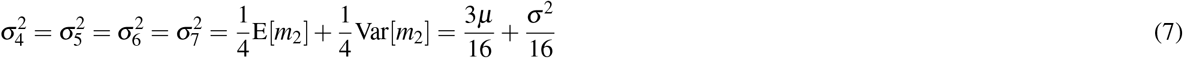

Indeed, solving the iterative equations (4) and (5), we get, for any cell *i* from generation *g* such that 2^g^ ≥ *i* > 2^*g*+1^,

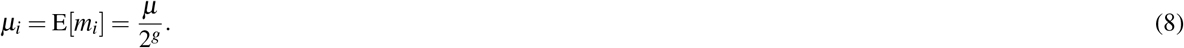

For the variance of *m_i_*, given *i* such that 2^g^ ≥ *i* > 2^*g*+1^, we get a somewhat more complex expression

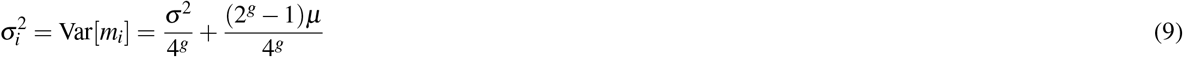

For *μ*_i_ and *σ*_i_ respectively in (8) and (9), since *i* is constrained by 2^g^ ≥ *i* > 2^*g*+1^, we can effectively substitute the cell label subscript *i* with the cell generation subscript *g*, such that for any integer value of *i*, the inequality 2^g^ ≥ *i* > 2^*g*+1^ is satisfied

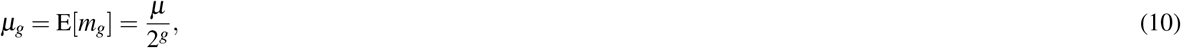

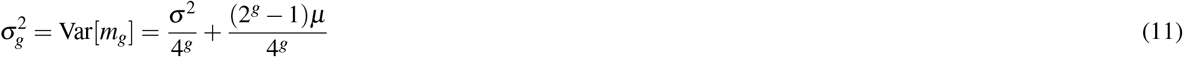

#### 3.1.2 Asymmetric case

If we assume 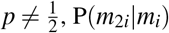 and P(*m*_2*i*+1_|*_i_*) have averages

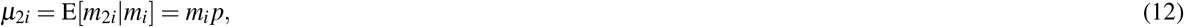

and

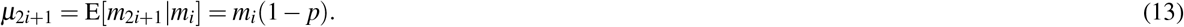

Similarly, for the variance, one finds

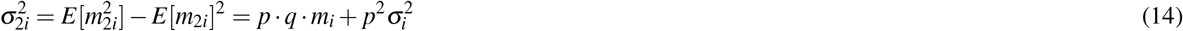

and

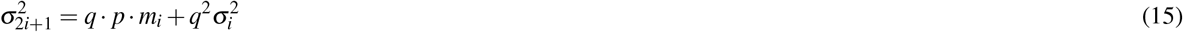

where *q* = 1 – *p*.

As the cell population grows, in an asymmetric division scenario, the different branches of daughter cells form subpopulations with distinct statistical expressions (see Fig. 3a). From these expressions, we understand how the mean and the variance of each branch in the lineage tree is linked to the mean and variance of the distribution of the initial progenitor cells population.

In contrast to the symmetric case, in which the two branches of daughter cells originated by the same sub-population of mother cells form identical distributions, in the asymmetrical case, both the means and the variances are different. In terms of generations, starting with an initial population of progenitor cells at generation 0 with a localized distribution, we end up with two distributions at generation one, four at *g* = 2, and 2^*g*^ at generation *g* (see Figure 3b-c). In general, our capability to distinguish the different sub-populations of each generation depends on the distribution parameters. In our case, the branch sub-population variance is too large to distinguish in the scatter plot the single generation branches Figure 4b). Thus, to compare the model with the experimental data, for which we can only distinguish the distribution for each generation, we need to compute the mean and the variance of the population generation, given by the superposition of all the lineage branches in the generation. Consequently, we want to compute for each generation

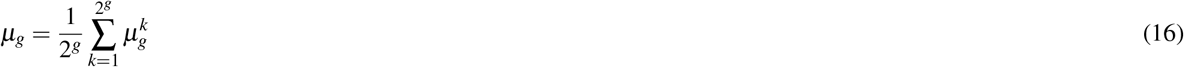

where 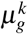 is the mean of the *k*-th sub-population of generation *g* and we assume that in each generation all the cells that replicate give rise to two daughters.

Knowing the initial population mean value, *μ*, equation (16) can be solved (see SI for more details) as

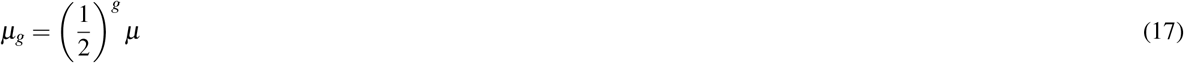

**Figure 4.**
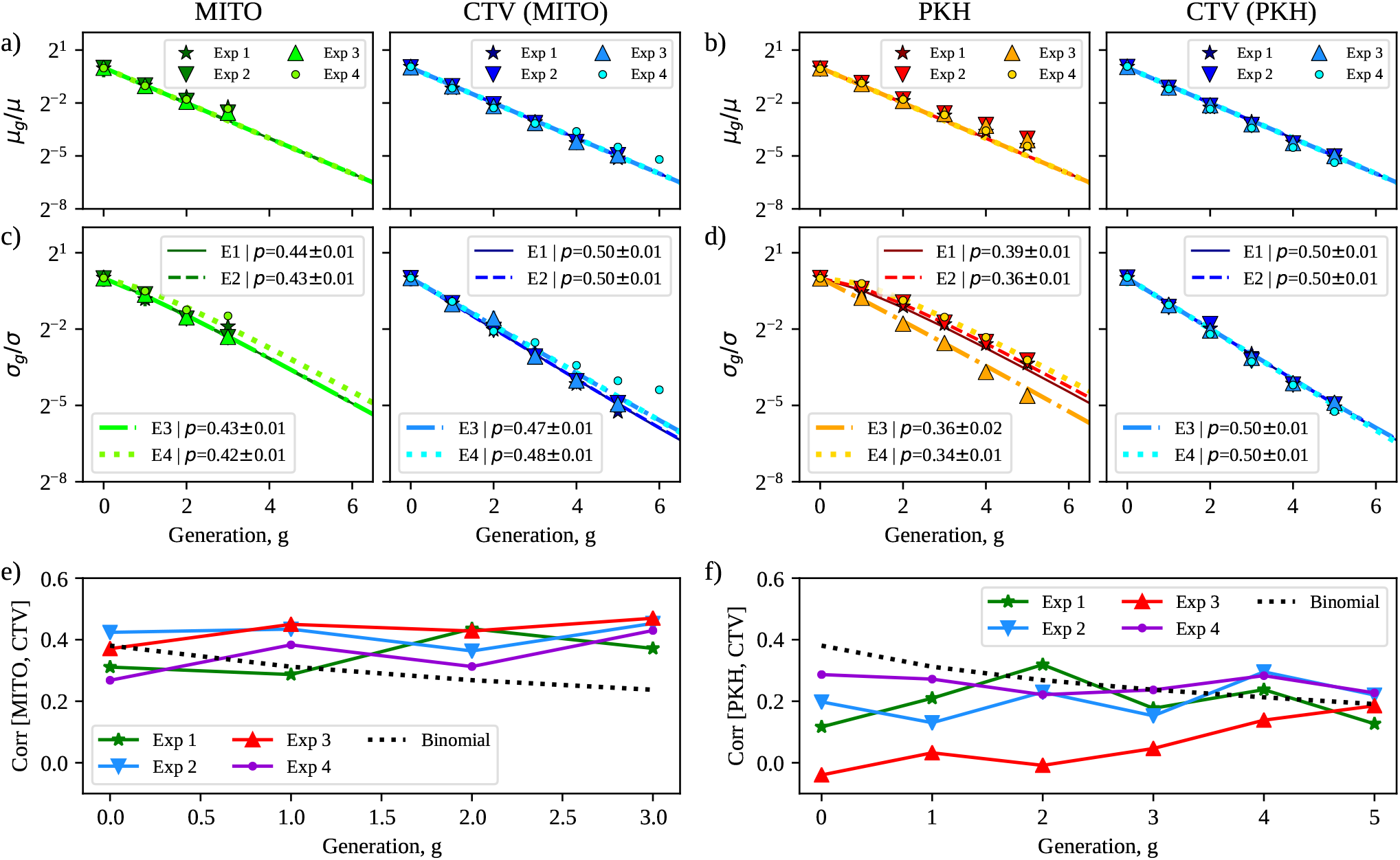
Model vs experiments. **a,b**) From left to right, the mean intensity of the three markers (MITO, CTV, and PKH) as a function of the generation, g, normalized over the mean intensity of the initial population. Dots correspond to experimental values obtained from the GM procedure while lines are the solutions of Eq. 17. CTV data has been analyzed twice, one time associated with MITO in the GM fitting and another with PKH. The results are very consistent. **c,d**) Same as in a) but with the variance of the intensity of each marker. Theoretical predictions (lines) are obtained from Eq. 19. **e**) Pearson correlation between the intensities of MITO and CTV markers as a function of the generations. The grey dotted line represents the predictions of an uncorrelated binomial partitioning, starting with a correlated initial population. **f**) Same as in c) but with the correlation between PKH and CTV markers.

It is interesting to note that the dependence of *p* (the coefficient of the binomial partitioning) cancels out because we are considering the mean of the mean values.

Consequently, to estimate *p* we must compute the variance. In the case of the variance, we want the variance of a distribution that is given by the mixture of the distributions of all the daughter cells sub-populations. It can be demonstrated that the total variance is given by

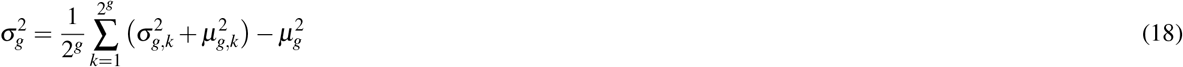

which after some computations, reported in the SI, yields

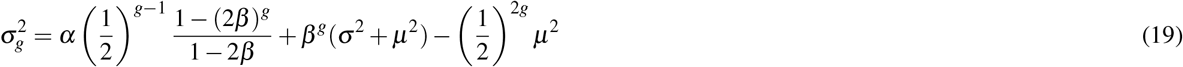

with *α* = *pqμ* and 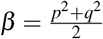. Figure 3d,e shows the behaviors of the mean fluorescence and its fluctuations as a function of the generations for different values of *p*. One can see that the trends are in line with those reported in Figure 2c,d. In the next section, we show that the proposed model quantitatively explains the observations.

## 4 Results

Firstly, we notice that according to the binomial partitioning model the mean value of the marker intensities at each generation is expected to depend only on the mean of the initial population, *μ*, and on the generation, *g* (see Eq. 17). Moreover, the *μ* dependence on *g* takes the form of a multiplicative constant, so that if we look at the mean intensity divided by the initial intensity as a function of the generation, we have a parameter-free way to confirm that the binomial model assumptions are at least partially correct. Figure 4a-b displays the mean fluorescence intensity of the three fluorescent dyes as a function of the generations divided by the mean intensity at generation 0.

We performed both a set of experiments with two-color stainings at a time (CTV-MITO and CTV-PKH) and another with three-color stainings (CTV-MITO-PKH). Working with two markers at a time ensured that during the sorting procedure a sufficiently large sample was collected. In fact, as for single marker flow cytometry to ensure peak detection, we must sort cells with a narrow gate centered at the dye intensity distribution mode. Consequently, the gate sorts a fraction, *f* of the total population. In a multicolor experiment, if we assume that each marker is gated so that a fraction *f* of cells is sorted for each color, in total we sort a fraction *f^d^* where *d* is the number of markers used. Thus, to achieve the same starting sample of a two-color experiment in a three-color experiment we have to prepare to sort a much larger population of cells than for a two-color experiment. Furthermore, due to the limitation of the sorter that we used with two markers, we set the collection gate and sorted all the stained cells in one sorting procedure(see Fig. 1b). While for a three-marker experiment we had to implement a subsequent sorting protocol with gates defined on the third-marker. Moreover, the two-color analysis allowed us to verify the consistency of the data because the CTV marker was present both in the two-color and three-color experiments.

In the Gaussian Mixture clustering procedure, we analyzed two colors at the time, Panel 4a shows how the mean changes when clustering was made on CTV vs MITO data, while Panel 4b shows the mean values for each generation, considering CTV and PKH intensities together. The CTV behaves identically in the two cases. Experimental data points are displayed with different markers, one kind for each experiment. Meanwhile, lines show the predictions of the asymmetrical binomial model in Eq. 17. Indeed, the intensity diminishes by a factor of two passing from one generation to the next for all four experiments, in perfect agreement with the binomial predictions.

However, considering the mean intensity of each generation does not allow us to evaluate if the partitioning process is symmetrical or not. In fact, as we can see in Eq. 17 and Figure 3d, the resulting mean intensity is unaffected by the partition fraction, p. Since the mean alone does not provide other information about the partition mechanism, we move to consider the variance. The variance analytical expression is given in Eq. 19, and it explicitly depends on *p*. Finally, we note that if we were able to distinguish the single sub-populations, forming the distribution of each generation (see Figure 3c), then we could study only the mean to fully characterize the partition mechanism. This can be seen in Eq. 12.

Next, we move on to analyze the fluctuations of the fluorescence signals, which are expected to inform us of the partitioning mechanism. In particular, Figure 4c,d displays the square root of the variance of the fluorescence divided by the square root of the variance at generation 0. Once more, data points are plotted in function of the generation, and with different markers for each one of the four reported experiments. Furthermore, lines in the same figure represent the results of the best fits using the theoretical expression for the variance of Eq. 19. The expression depends on three parameters, i.e. the mean intensity at generation zero, *μ*, the initial variance, *σ*^2^, and the partition probability, *p*. Fixing the *μ*s to those found in the analysis of the mean intensities, we are left with two free parameters. In particular, the partition probability, *p*, is of particular interest since it gives a measure of the asymmetry. Clearly, with *p* = 0.5 we get a completely symmetric partitioning process, and for *p* → 0, we get an asymmetric or biased partitioning process.

As for the mean, the variances are also found in excellent agreement with the predictions of the asymmetric binomial partitioning model for all the three cellular constituents we analyzed, although with different partition probabilities. In particular, cytoplasm appears to undergo a symmetrical partition, with a *p* of ~ 0.50 both when measured in couple with mitochondria (Figure 4c) and membrane proteins (Figure 4d). Since the amount of cytoplasm is expected to correlate with the cell volume, its symmetrical partitioning is informative on the population size distribution and in accordance with what is known on Jurkat cell division.

On the other hand, mitochondria and membrane proteins are both unevenly partitioned, i.e. they have a *p* significantly lower than 0.5. In particular, mitochondria fluorescence variance is well reproduced with *p* ~ 0.43, as shown in Figure 4c. This asymmetry in mitochondria partitioning has been observed in yeast cells and mammalian epithelial stemlike immortalized cells. Furthermore, it was connected to the anti-aging defense of the population^27,44–47^. Similarly, a partitioning probability of ~ 0.36 gives the best description of the fluorescence variance associated with the PKH dye (Figure 4d). In this case, the biased segregation may be linked to aggregates of membrane proteins (rafts), that T-cells form when activated, as well from endocytosed florescence. Indeed, Jurkat cells are known to form clusters of membrane proteins. Those microdomains are required for protein-protein interaction and, during activation, are implicated in T cells signaling^48^. To verify that the membrane signal was due in some cases to clusters of aggregation, we acquired confocal microscopy images at the starting point. Indeed, clusters of signals at the membrane level for the PKH fluorescence were evident in many of the analyzed cells. Furthermore, as described by Katajisto *et al*.^27^ endocytosed PKH formed distinct cytoplasmic puncta.

We want to stress that we were able to characterize mitochondria and membrane partitioning because they were associated with the cytoplasmic symmetrical marker. In fact, we show that notably symmetrical inherited dyes also have an additional practical interest because they can be used to support peak detection of noisier markers^40^. Indeed, while CTV is inherited in a highly symmetrical fashion in Jurkat cells and shows well-resolved division peaks, both the other dyes tend to partition asymmetrically and to lose fluorescence during the culturing conditions. These two effects add together to broaden the fluorescence distributions. As one can see in the examples from Figure 1e and g, where the single marker marginal distributions and the two marker distributions are shown together, the two generations are hardly distinguishable if we look at the PKH intensity alone, while they are well separated with respect to the CTV marker. Thus, the strategy of using different dyes enables us to reduce the variation in the resolution peak for each generation and also to analyze correlations among different markers. In our case, the simultaneous staining of mitochondria and cytoplasm, and that of membrane and cytoplasm, enables us to investigate the correlations between their partitioning mechanisms.

Specifically, we measured the Pearson correlation coefficient between the intensity of MITO and CTV and between PKH and CTV intensities in different generations. As one can see from Figure 4d,e, the fluorescences of both couples tend to present an initial correlation higher than zero. The correlation of mitochondria and cytoplasm (MITO-CTV) is ≃ 0.35, while that of membrane proteins and cytoplasm is *ρ* ~ 0.20. During proliferation, both correlations are preserved from one generation to the next for all experiments where the initial distributions are correlated. Interestingly, the only experiment that starts with an uncorrelated population (Exp 3 in Figure 4e) shows a progressive increase of correlation which reaches the final value of the other experiments. Notably, the observed behavior of correlations is not compatible with uncorrelated two-variable binomial statistics. In fact, in this case, the expectation is a correlation that decreases along over the generations even if the population of cells starts with a correlated distribution of stained compounds (the grey dotted line in panels d and e).

## 5 Discussion

Phenotypic heterogeneity has gained much attention in recent years as novel high-throughput experiments became able to study cell populations in greater detail. In particular, variability of non-genetic origin has been found to play a dominant contribution of cancer heterogeneity and to act as a driving force for tumor evolution^49,50^. While such variability is found in many biological aspects, its origin is often ascribed to stochasticity at the level of gene expression and its regulation. However, fluctuations also arise during cell division when molecules are partitioned stochastically between the two daughter cells^2^. Disentangling the two contributions is quite difficult, and it has been suggested that often experimental outcomes may have been misinterpreted^15^. In fact, experiments often look at phenotypic traits to infer the effects of partitioning errors, even though those indirect observables are influenced by homeostatic pressure and thus by regulation. To root out these problems, we devised an experimental and computational apparatus to follow the proliferation of cell populations and directly measure the partitioning of different cellular elements at the same time by staining cellular components with fluorescent markers at the beginning, upon cell sorting, and then measuring the dynamics of the partitioning process. This assures us to limit expression effects.

Partitioning of cellular compounds is expected to depend on the particular types of compounds. Intuitively, those compounds that are present in huge numbers and are uniformly distributed across the whole cell could follow a random segregation mechanism. Meanwhile, both the replication and partitioning of nuclear chromosomes must be stringently controlled, with each chromosome being replicated exactly once per cell cycle and one copy transmitted to each daughter cell. To ensure perfect hereditary continuity of all the genes, stringent control is necessary to avoid the loss of essential genes in some cells. In fact, errors in the duplication and partitioning of genes or entire chromosomes are the cause of important genetic disorders, such as trisomy 21^1^. The scenario is less clear for cell organelles and internal compounds. Many RNAs, proteins, and organelles are present in numbers low enough that segregation of individual copies causes large fluctuations at cell division. Moreover, the presence of biases in the partitioning of some compounds can produce a systematic difference in the daughters, which fosters cell differentiation^51^. Even symmetrically dividing cells can then produce daughters with very different compositions by chance. Depending on the rates with which compounds are restored to their physiological levels, the single-cell can generate variability. Most interestingly, as much as partitioning noise appears to take part in several cell division processes, cell populations are characterized by homeostasis that tends to contrasts with noise^52, 53^. Consequently, partitioning noise and other stochastic factors must cooperate to coordinate the population in a collective homeostatic state, similarly to other stochastic system present emergent collectively coordinate^54–58^

Mitochondria and endosomes are known to exhibit asymmetrical partitioning in yeast^44, 45, 59–62^ and in asymmetrically dividing cells, like mammalian epithelial stem-like immortalized cells^27, 63^. This asymmetrical mitochondria partitioning enables cells to defend themselves from aging because it protects the cell progeny from accumulating misfolded proteins engulfed in the mitochondria^27,44–46^.

Similarly, biased segregation of membrane proteins in bacteria has been linked to the formation of protein raft which migrates differently to the older pole in the mother cell with respect to the novel one in the daughter cell, thus influencing the aging of the overall population^64^. Moreover, Bergmiller *et al*.^12^ found that biased binomial segregation helps bacterial populations to face antibiotic treatments, suggesting partitioning as a drug resistance enhancer. Whether similar mechanisms take place also in tumoral cells is still under scrutiny.

In the present work, we investigate the partitioning of three important cellular components during the proliferation of cancer T-cell (Jurkat) populations. In particular, we use three distinct fluorescent markers to label mitochondria, membrane proteins, and cytoplasm and quantified for the fist time, to our knowledge, both the degree of segregation asymmetry of each element between the two daughter cells and the correlations in the partitioning of different kinds of compounds. Indeed, comparing the fluorescence mean intensities and their fluctuations against the predictions of a biased binomial model, we found that mitochondria and double layer lipid membrane partitions asymmetrically while cytoplasm partitions symmetrically. Nevertheless, it is important to consider that in comparison, to mammalian epithelial stem-like immortalized cells that asymmetrically partition more than five times the old mitochondria in the mother cell, for Jurkat cell the bias is much smaller with a biased probability of around 0.43.

This indicates that in Jurkat cells asymmetrical segregation is present even if in a weaker form in comparison to what is observed for stem-like asymmetrically dividing cells. Second, it shows how our segregation asymmetry measurement method takes advantage of the large number of cells that can be processed in a flow cytometry experiment to improve the experimental resolution.

Begum *et al*.^40^ shows that succinimidyl fluorescent dyes can be used to track cell proliferation for a maximum of 72 h, while Filby *et al*.^65^ demonstrates that lipophilic dyes peaks are already hard to distinguish at 48 h, concluding that lipophilic dyes are weak proliferation trackers. Our multicolor approach allows us to resolve the peaks formed by the different generations up to 72 h for the lipophilic dye PHK, as for the CTV. This is because with the use of multicolor flow cytometry, we were able to merge the information gathered from CTV on the analysis of the PKH. Finally, we can measure and analyze partitioning error across different cell generations, and analyze how the covariance of the partitioning compounds changes. This allows us to show the gradual weakening of the PKH lipophilic dye peak resolution is the result of the partitioning noise that broadens the peaks. We also used the MitoTracker dye together with CTV and with both CTV and PKH, this allowed us to track how progenitor mitochondria are inherited by the progeny.

In conclusion, in contrast to a single-color flow cytometry live cell experiment that enables tracking cell proliferation of only one dye at a time^65^, in our multicolor experiment, we were able to simultaneously follow more than one live cell marker and quantify their partitioning during cellular proliferation up to more generations than one can do using only one marker at a time.

Finally, we want to stress that our apparatus can be easily applied to different cell lines and different kinds of compounds providing a powerful tool for understanding partitioning-driven heterogeneity. Moreover, our method is not based on transfection, and therefore in the future, it may be used in primary cell cultures. In perspective, this could be particularly relevant in the case of tumor microenvironment diversity, where comprehension of the high cell heterogeneity could pave the way to novel and more targeted therapies.

## Materials and Methods

### Cell culture

E6.1 Jurkat cells (kindly provided by Dr. Nadia Peragine, Department of Cellular Biotechnologies and Hematology, ‘Sapienza’ University) were used as a cell model for proliferation study and maintained in RPMI-1640 complete culture media containing 10% FBS, penicillin/streptomycin plus glutamine at 37*°* C in 5% CO2. Cells were passaged 3 times prior to amplification for the experiment. Cells were then harvested, counted, and washed twice in serum-free solutions and re-suspended in pre-warmed PBS for further staining.

### Cells fluorescent dye labeling

To track cell proliferation by dye dilution establishing the daughter progeny of a completed cell cycle, cells were stained with PKH26, CellTraceTM Violet (CTV), and MitoTracker^®^ Green FM according to the manufacturer’s instruction. The use of three different cell markers, respectively cell membrane, cytoplasm, and mitochondria, allowed for a multi-variable proliferation analysis. Fixable Viability Stain 780 (FVS780) was used in each experiment to determine cell viability and verify low dye toxicity. The first staining was performed with PKH26, a red fluorescent dye used for Cell Tracking Lipophilic Membrane Dyes (MINI26-1KT, Sigma, St. Louis, USA). Briefly, 20 × 10^6^ cells were collected, washed once in PBS, and resuspended in 500ul of Diluent C for lipophilic dyes. The cell suspension was then added to the same volume of 2x Dye Solution (4uM) in Diluent C by adding 4ul of the PKH26, mixed gently, and kept for 2 min in gentle agitation using vortex at room temperature (RT). Immediately after, the cellular suspension is blocked with an equal volume of complete FBS, added additional complete media to fill the tube, and centrifuged. Cells were washed again once in PBS prior to the following staining. The CTV dye (C34557, Life Technologies, Paisley, UK), typically used to monitor multiple cell generations, was performed according to the manufacturer’s instruction, diluting the CTV 1/1000 in 1ml of PBS for 20 min at RT. Afterward, complete media was added to the cell suspension for additional 5 min incubation before the final washing in PBS. MitoTracker^®^ Green FM (M7514, Molecular Probes, Eugene, USA), green-fluorescent mitochondrial stain, was used at the concentration of 20nM (stock 20um in DMSO) in PBS for 30 min in a water bath at 37*°*C, mixing every 10min. Cells were then washed once in PBS. The live/dead staining, Fixable Viability Stain 780 (565388, BD Biosciences, San Jose, USA) was usually performed last, 1/1000 in PBS for 10min at RT. For each experiment, a new aliquot of Viability marker was used.

### Cell sorting

Jurkat cells labeled with dyes were sorted using a FACSAriaIII (Becton Dickinson, BD Biosciences, USA) equipped with Near UV 375nm, 488nm, 561nm, and 633nm lasers and FACSDiva software (BD Biosciences version 6.1.3). Data were analyzed using a FlowJo software (Tree Star, version 9.3.2). Briefly, cells were first gated on live cells, viability dye exclusion, followed by doublets exclusion with morphology parameters, both side and forward scatter, area versus width (A versus W). The unstained sample was used to set the background fluorescence for each channel. For each fluorochrome, a sorting gate was set around the max peak of fluorescence of the dye distribution (see ^65^). In this way, the collected cells were enriched for the highest fluorescence intensity for all the markers used according to the experimental setup: PKH26, CTV, and/or Mito Green. Following isolation, an aliquot of the sorted cells was analyzed at the same instrument to determine the post sorting purity and population width, resulting in an enrichment > 99 % for each sample (see Fig S1 in the Supporting Information).

### Cell culture for dye dilution assessment

Sorted cell population was seeded in 6 wells plates (BD Falcon) at 1×10*6 cells/well and kept in culture up to 72 hours. To monitor multiple cell division, an aliquot of the cells in culture was analyzed every 18, 24, 36, 48, 60, and 72 hours for the expression of the different markers by LSRFortessa flow cytometer. For setting the time zero of the kinetic, prior culturing, a tiny aliquot of the collected cells was analyzed immediately after sorting at the flow cytometer. The unstained sample was used to set the background fluorescence as described above. Every time that an aliquot of cells was collected for analysis, a same volume of fresh media was replaced to the culture.

### Expectation-Maximization and the Gaussian Mixture Model

We used the Expectation-Maximization (EM) algorithm to detect the clusters in Gaussian Mixture Models^66^. The EM algorithm is composed of two steps the Expectation (E) step and the Maximization (M) step. In the E-step, for each data point **f**, we used our current guess of *π_g_*, *μ_g_* and σ*_g_*, to estimate the posterior probability that each cell belongs to generation *g* given that its fluorescence intensity measure as **f**, *γ_g_* = P(*g*|**f**). In the M-step, we use the fact that the gradient of the log-likelihood of *p*(**f_i_**) for *π_g_*, *μ_g_* and *σ_g_* can be computed. Consequently, the expression of the optimal value of *π_g_*, *μ_g_* and *σ_g_* is dependent on *γ_g_*. It is shown that under, certain smoothness condition the iterative computation of E-stem and M-step leads us to the locally-optimal estimate of the parameters *π_g_*, *μ_g_* and *σ_g_*, and returns the posterior probability *γ_g_* which weights how much each point belongs to one of the clusters. Here, we used this model to perform cluster analysis and detect the peaks which correspond to different generations. Then, from these clusters we estimated *π_g_*, E[*f_g_*], and Var[*f_g_*].

### Mixture of Gaussian Clustering

To estimate to which generation the detected cell belongs, we used the mixture of Gaussian method. We always used all two fluorescence measures for the clustering method, for the experiments with two dyes (CTV-PKH and CTV-MITO). We used all the fluorescence measures for the clustering method, for the experiments with three dyes (CTV, PKH, and MITO), except in the samples where the MITO after being diluted in the successive generation got very close to the background fluorescence and added too much noise to the clustering method biasing the result. Fig.1d shows an example of how the mixture of the Gaussian method was able to cluster the data.

## Supporting information

Supplemental calculations and figures

## Author contributions statement

G.G. conceived research; G.R. contributed with additional ideas; G.P., R.M., and G.G. performed experiments; M.M., G.P., and G.G. analyzed data; M.M. and G.G. performed analytical calculations and statistical analysis; all authors analyzed results; all authors wrote and revised the paper.

## Acknowledgements

The authors are indebted with Andrea De Martino, Andrea Gamba, and Andrea Pagnani for fruitful and enjoyable discussions.

