## Supplemental calculations and figures for "Asymmetric Binomial Statistics Explains Organelle Partitioning Variance in Cancer Cell Proliferation"

### I. GENERAL BINOMIAL MODEL

Let us suppose that a population of cells is free to grow in a controlled environment and we marked some of the components of each cell with a fluorescent die (for example the mitochondria or the membrane proteins). Focusing on one compound species, we assume that at time zero each cell has $m_i$ marked compounds, in such a way that if we measure the whole population we get a marker distribution of mean $\mu$ and variance $\sigma^2$.

Each cell of the population grows and divides at a certain rate so that if we monitor the population at different times, we can follow the evolution of the marker intensity distribution. Since from each mother cell two daughters are generated at each duplication the marked compounds split between the two daughters with a certain ratio, which indeed is one of the parameters we want to characterize. Assuming a binomial partitioning of the compounds, we will have that the probability of finding a cell with $m_{2i}$ marked compounds at generation $g$, is given by:

$$P(m_{2i}|m_i) = \binom{m_i}{m_{2i}} p^{m_{2i}} q^{m_i - m_{2i}}$$

with $q = 1 - p$. In general, the expected value and the variance of such a distribution are given by $p \cdot m_i$ and $p \cdot q \cdot m_i$, respectively. Since we do not study a single cell, but a population of cells, we must deal with distributions, that evolves according to

$$P(m_{2i}) = \int P(m_{2i}|m_i)P(m_i)dm_i. \tag{1}$$

We now want to derive a closed-form equation of the expectation and the variance of $P(m_i)$. To derive the expectation value, we use the total expectation law, which is just a consequence of the double-integrals expression of the expectation value. Assuming that the population indexed with $2i$ is associated with $p$ and the one with $2i + 1$ is associated with $q = 1 - p$, we have that

$$\mu_{2i} = E[m_{2i}] = \int dm_{2i}P(m_{2i})m_{2i} = \int dm_i dm_{2i}P(m_{2i}|m_i)P(m_i)m_{2i} = E[m_i p] = \mu_i p \tag{2}$$

and similarly

$$\mu_{2i+1} = E[m_{2i+1}] = \int dm_{2i+1}P(m_{2i+1})m_{2i+1} = E[m_i q] = \mu_i(1 - p) \tag{3}$$

For the variance, one can use a similar approach

$$\sigma^2_{2i} = E[m^2_{2i}] - E[m_{2i}]^2 = E[E[m^2_{2i}|m_i]] - E[E[m_{2i}|m_i]]^2 = E[Var[m_{2i}|m_i]] + Var[E[m_{2i}|m_i]] = p \cdot q \cdot m_i + p^2\sigma^2_i \tag{4}$$

and

---

* The authors contributed equally to the present work.

$$\sigma^2_{2i+1} = q \cdot p \cdot m_i + q^2\sigma^2_i \tag{5}$$

Now, we know how the mean and the variance of each sub-population of daughter cells is linked to the mean and variance of the population of mother cells. To compare the model with the experimental data, we need to compute the mean and the variance of the whole population (given by the over position of all the daughter cell sub-populations). We want to compute for each generation

$$\mu_g = \frac{1}{2^g}\sum_{k=1}^{2^g}\mu^k_g \tag{6}$$

where we assume that in each generation all cells replicate giving rise to two daughter (if this is not the case, the $\frac{1}{2^g}$ term must be substituted with the relative abundance of each sub-population in the summation). Making use of Eq. 2 and 3, we can split the sum in Eq. 6 in two groups, the one with the sub-population of daughter that inherits a '$p$' fraction of compounds and the other that inherits a '$q$' fraction of compounds, i.e.

$$\mu_g = \frac{1}{2^g}\left[\sum_{k=1}^{2^{g-1}} p\mu^k_{g-1} + \sum_{k=1}^{2^{g-1}}(1-p)\mu^k_{g-1}\right] \tag{7}$$

which can be recast as

$$\mu_g = \frac{1}{2^g}\left[\frac{2^{g-1}}{2^{g-1}}\sum_{k=1}^{2^{g-1}}\mu^k_{g-1}\right] = \frac{1}{2}\mu_{g-1} \tag{8}$$

Knowing the initial population mean value, $\mu$, this equation can be solved recursively as

$$\mu_g = \left(\frac{1}{2}\right)^g \mu \tag{9}$$

It is interesting to note that the dependence of $p$ (the coefficient of the binomial partitioning) cancels out since we are considering the mean of the mean values.

To estimate the $p$ we must compute the variance. In the case of the variance, we want the variance of a distribution that is given by the mixture of the distribution of all the sub-populations of daughter cells. In this case, it can be demonstrated that the total variance is given by

$$\sigma^2_g = \frac{1}{2^g}\sum_{k=1}^{2^g}\left(\sigma^2_{g,k} + \mu^2_{g,k}\right) - \mu^2_g \tag{10}$$

To obtain a recursive expression also for the variance, we can recast the equation as

$$A_g = \sigma^2_g + \mu^2_g = \frac{1}{2^g}\sum_{k=1}^{2^g}\left(\sigma^2_{g,k} + \mu^2_{g,k}\right) = \frac{1}{2^g}\left[\sum_{k=1}^{2^{g-1}} {}^{(p)}\sigma^2_{g,k} + {}^{(p)}\mu^2_{g,k} + \sum_{k=1}^{2^{g-1}} {}^{(q)}\sigma^2_{g,k} + {}^{(q)}\mu^2_{g,k}\right] \tag{11}$$

where the label $^{(p)}$ (resp. $^{(q)}$) indicates that the daughter inherits a 'fraction' p (resp. q) of compounds. Making use of Eq. 2, 3, 4 and 4 we can recast Eq. 11 as

$$A_g = \frac{1}{2^g}\left[2pq\sum_{k=1}^{2^{g-1}}\mu_{g-1,k} + (p^2+q^2)\sum_{k=1}^{2^{g-1}}\left[\sigma^2_{g-1,k}\mu^2_{g-1,k}\right]\right] =$$

$$= \frac{1}{2^g}\left[\frac{2^{g-1}}{2^{g-1}}2pq\sum_{k=1}^{2^{g-1}}\mu_{g-1,k} + (p^2+q^2)\frac{2^{g-1}}{2^{g-1}}\sum_{k=1}^{2^{g-1}}\left[\sigma^2_{g-1,k}\mu^2_{g-1,k}\right]\right] = pq\mu_{g-1} + \frac{p^2+q^2}{2}A_{g-1} \tag{12}$$

Making of of Eq. 8 we can recast the equation for the variance as

$$A_g = \alpha \frac{1}{2}^{g-1} + \beta A_{g-1} \tag{13}$$

with $\alpha = pq\mu$ and $\beta = \frac{p^2+q^2}{2}$.

The previous equation can be expanded and rewritten as

$$A_g = \alpha \left(\frac{1}{2}\right)^{g-1} \sum_{k=0}^{g-1} (2\beta)^k + \beta^g A_0 = \alpha \left(\frac{1}{2}\right)^{g-1} \frac{1-(2\beta)^g}{1-2\beta} + \beta^g A_0 \tag{14}$$

and finally one obtains

$$\sigma_g^2 = \alpha \left(\frac{1}{2}\right)^{g-1} \frac{1-(2\beta)^g}{1-2\beta} + \beta^g(\sigma^2 + \mu^2) - \left(\frac{1}{2}\right)^{2g} \mu^2 \tag{15}$$

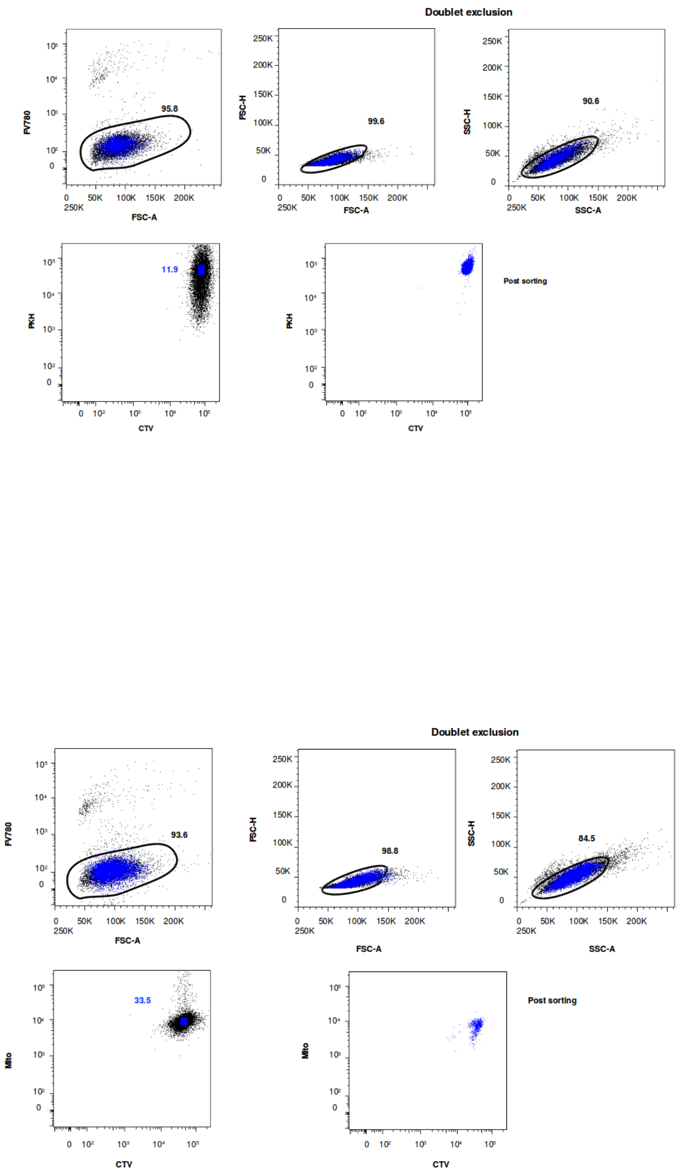


FIG. 1: **Gate strategy for isolation of PKH/CTV/Mito positive Jurkat cells.**
Representative dot plots of Jurkat cells showing the gating strategy for isolation of PKH/CTV/Mito highly positive cells. Live cells first gated based on forward scatter area parameter (FSC-A and FV780 negative cells) were then selected for doublets exclusion (Heights, H versus Area, A for both FSC and side scatter; upper panels) an¡d finally collected based on PKH and CTV or Mito and CTV highest expression level (left lower panels). Double positive cells sorting efficiency are shown in the right lower panel. To show the sorted cells (blue) back gating strategy is used in all plots to identify the cell population and to confirm the gating strategy. .
